# Production of high-affinity glycosylated anti-mouse conjugated nanobodies in *Pichia pastoris*

**DOI:** 10.1101/2025.07.25.666866

**Authors:** Sofía Orioli, Javier Santos, Lorena I. Ibañez, Cecilia D’Alessio

## Abstract

Nanobodies (NBs) are small antibody fragments derived from camelid heavy-chain antibodies which represent the minimal functional domain capable of antigen recognition and binding. NB are ten times smaller than conventional antibodies, exhibit a compact structure and high stability, making them ideal for recombinant production. The eukaryotic unicellular system *Pichia pastoris* provides multiple advantages for protein expression, including the ability to perform several eukaryotic posttranslational modifications. In this work, we engineered a modular plasmid sequence that, through specific restriction enzyme cuts and ligations, codes the expression of a secreted anti-mouse *kappa* chain NB fused with various accessory peptides in *P. pastoris*. This system enables the incorporation of a plastic binding sequence, a histidine tag (Hisx6) for purification, the horseradish peroxidase (HRP) enzyme for chemiluminescence detection, or the biotinylatable AviTag sequence, in multiple combinations. We successfully expressed and purified anti-*kappa* NBs fused to a Hisx6-tag (κNB) and to HRP -Hisx6-tag (κNB-HRP), with subsequent structural and functional characterization revealing high affinity for mouse immunoglobulins. The κNB-*kappa* light chain domain complex was modeled showing a fitted surface interaction of CDR3 domain. The position of a glycan in the complex was modeled predicting that glycan addition would not affect the interaction surface. Accordingly, no functional differences were observed in κNB after deglycosylation, indicating that high mannose glycan addition has not interfered with its binding capability. Moreover, glycosylated κNB fused to HRP was expressed with retained HRP activity, and proved to be functional as a secondary antibody, demonstrating the system’s versatility in producing NBs and conjugated NBs with posttraslational modification that may be required for diverse biotechnological applications.

## INTRODUCTION

Nanobodies (NBs) are the smallest antibody fragments capable of recognizing and binding antigens with high specificity and affinity^[1,2]^. NBs consist solely of the ∼15 kDa heavy-chain variable region (V_HH_) of homodimeric heavy-chain antibodies discovered in camelids and sharks in the 1990s^[3–5]^ **(Figure 1A)**. Due to their small size, NBs possess high chemical, thermal, and structural stability; high solubility; and the ability to bind to epitopes that are difficult to access for larger conventional antibodies^[2,6]^. The aforementioned properties, along with their simple structure, makes NBs excellent candidates for recombinant expression in microorganisms.

**Figure 1:**
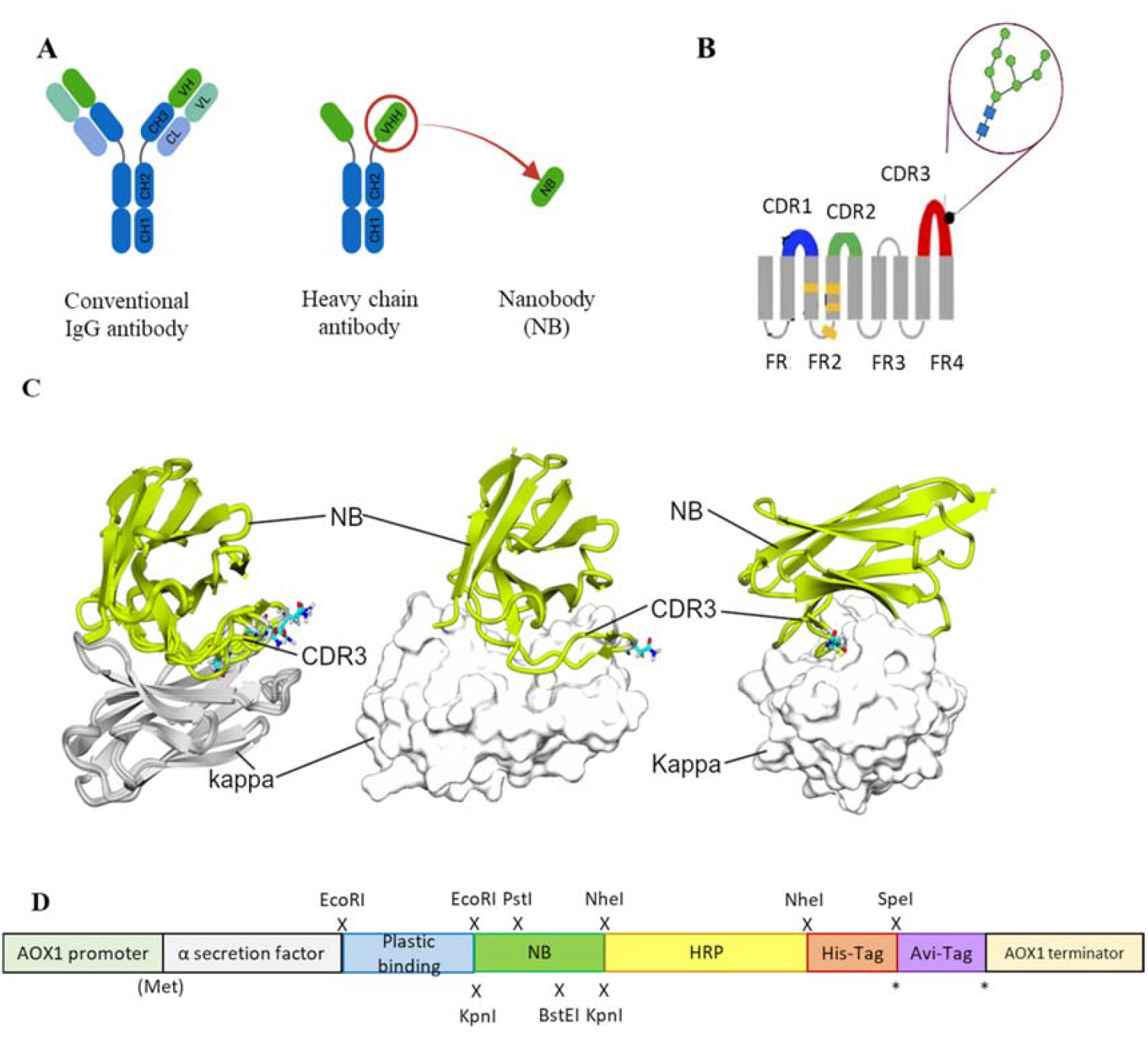
(A) Schematic representation of different antibody formats. On the left, a typical mammalian IgG antibody; in the center, a camelid heavy-chain IgG; and on the right, a VHH or NB. The acronyms stand for: CH, heavy-chain constant domain; CL, light-chain constant domain; VH, heavy-chain variable domain; and VL, light-chain variable domain (Created in https://BioRender.com). (**B**) NB diagram showing the positions of CDR domains: CDR1 (blue), CDR2 (green) and CDR3 (red) and the predicted position of a single *N*-glycan (Figure adapted from Mitchell & Colwell,2018^[14]^). **(C)** Interaction κNB and the mouse *kappa* light chain. In the left panel, a superimposition of five AlphaFold3 models is shown interacting with mouse *kappa* light chain (gray, *kappa*; green, κNB). The Asn107 residues in each model are depicted in sticks. In the central panel, only one model is presented, with the *kappa* chain shown in surface representation, similar to the right panel, where the model’s orientation was changed to highlight the extensive interaction of CDR3. (**D**) Modular insert designed for the expression of a NB fused to different accessory peptides/ proteins in the pPICZαA vector. From 5’: AOX1 promoter to drive expression, *Sacharomyces cerevisiae* Mating Factor α secretion signal, Plastic Binding tag, exchangeable NB, Horse Radish Peroxidase, Hisx6 tag, AviTag sequence and at the 3’ end the AOX1 transcription terminator sequence. The restriction enzyme cleavage sites introduced are indicated by X, designed to generate different NB variants. The asterisk * shows the location of alternative translation termination stop codons. The legend (Met) indicates the translation initiation.

*Escherichia coli* is the most commonly used host for NB production in laboratories. However, both cytoplasmic expression and periplasmic secretion present significant challenges. In the periplasm, the limited compartment size can become overcrowded when NBs are highly expressed, overwhelming the chaperone system and impairing proper protein folding. In the cytoplasm, the reducing environment hinders disulfide bond formation, often resulting in NB aggregation into inclusion bodies, which cannot be always successfully refolded^[1,7]^. Moreover, generating a conjugated NB whose partner is not expressed efficiently in bacterial systems would be a bottleneck^[8,9]^. The methylotrophic yeast *Pichia pastoris* has emerged as a popular eukaryotic expression system as it can perform post-translational modifications such as *N*- and *O*-glycosylation, disulfide-bond formation, and protein folding in the secretory pathway^[10]^, while remaining easy to genetically modify and culture. Its highly efficient secretory pathway facilitates protein purification from culture supernatants when expressed proteins are fused to secretion peptides. Although several promoters have been used in expression vectors, the AOX1 (alcohol oxidase 1) promoter-inducible by the addition of methanol-is still very popular as it enables high, stable, and controlled expression of recombinant proteins^[10–12]^.

Given that ∼99% of commercial mouse antibodies contain the *kappa* light chain ^[13]^, an anti-*kappa* NB (κNB) is highly valuable for various applications like immunoassays, Western Blots and ELISA assays. Generating in-house conjugated variants of anti-mouse *kappa* chain NBs from a single DNA sequence will enable diverse applications in research and diagnostics.

In this work, we expressed functional anti-mouse *kappa* chain NBs in the methylotrophic yeast *P. pastoris* using a single modular DNA sequence that through differential cuts and ligations allows obtaining the NBs fused to diverse accessory peptides that ease detection and purification. We also showed that glycosylation not only did not affect the binding capability of κNB, but also allowed its fusion to a conjugated enzyme (HRP) whose expression benefits from glycosylation. Glycans may be removed for homogeneity purposes after purification without loss of functionality.

## RESULTS

### Interaction of κNB with mouse immunoglobulin *kappa* light chain

AlphaFold3 was used to predict the structure of the complex between the κNB and the mouse *kappa* variable domain (BBA57852.1). It is worth mentioning that the κNB sequence contains one consensus sequence for *N*-glycosylation (N-X-S/T, where X can be any amino acid except P). This sequon is located between amino acids 107-109 in the κNB, corresponding to the CDR3 **(Figure 1B)**. The predicted model for the κNB-*kappa* complex yielded ipTM and pTM values of 0.71 and 0.78, respectively, indicating that the structure could be similar to the actual structure. The most inaccurate region of the model was the CDR3 stretch of the κNB, which, paradoxically, was involved in the direct interaction between κNB and the *kappa* domain. Interestingly, the NXS glycosylation site located in the CDR3 region was exposed to the solvent in 4 of 5 models, suggesting that if the CDR3 adopted these conformations, the glycan would not disrupt the interaction between the κNB and *kappa* domains. It is worth noting that the CDR3 region creates a large contact area between both domains, which could contribute to the complex’s high stability **(Figure 1C)**.

### Modular plasmid design for the expression of κNBs fused to different accessory peptides

A unique pPICZαA vector was designed for the expression in *P. pastoris* of an anti-κ NB in fusion to different tags and/or accessory peptides which provide a variety of features, such as a plastic binding sequence, the HRP, the biotinylatable AviTag sequence and a 6-histidine tag. Each coding sequence was flanked by the sequences of different restriction enzymes to allow the specific removal of each individual peptide, after which the plasmid can be re-ligated and different nanobody variants expressed (**Figure 1D**). All sequences were preceded by an alpha factor pre-pro-sequence, used to secrete the expressed protein to the yeast culture supernatant. Then, the modular vector contained the following coding sequences: a plastic binding peptide (flanked by EcoRI sites); the anti-mouse κNB (flanked by KpnI, PstI and BstEI sites); the HRP (flanked by NheI sites); a Hisx6 tag (after which there is a stop codon followed by a SpeI site); and lastly a biotinylatable AviTag sequence, also followed by a stop codon. As restriction with NheI and SpeI result in compatible cohesive ends, restriction with both enzymes allows for the removal of both HRP and the Hisx6 tag, and the expression of the NB fused to the AviTag sequence instead of HRP as a means for detection. Similarly, restriction with EcoRI followed by re-ligation would remove the possibility to bind the protein to plastic leaving only the Hisx6 tag to facilitate purification. The new modular vector was named pPICZαA-Plastic-κNB-HRP-Hisx6-AviTag (henceforth referred as “Modular vector”).

Restriction reactions, re-ligations and amplifications were carried out to obtain different variations of the plasmid. The HRP detection sequence was removed with NheI and the plastic binding sequence with EcoRI, resulting in a plasmid for the expression of κNB fused only to the Hisx6 tag to test the system (κNB). Simultaneously, a vector for the expression of the κNB fused to HRP and the Hisx6 tag was obtained by restriction only with EcoRI (κNB-HRP). All new plasmids were verified by sequencing and integrated in *P. pastoris* X-33 genome.

### Expression and characterization of κNB expressed in *Pichia pastoris*

Expression of κNB was carried out in 200 mL cultures of *P. pastoris* strain X-33 transformed with the plasmid digested with NheI and EcoRI and re-ligated. Inductions were performed with 1% methanol and every 24 hours, aliquots were taken, and proteins were precipitated with 15% trichloroacetic acid. The presence of the κNB was confirmed by Western Blot using a mouse anti-His primary antibody followed by an anti-mouse-HRP secondary antibody (**Figure 2A**). A progressive increase in protein quantity over time was observed with a maximum upon 72 hours of methanol induction. κNB has an expected molecular mass of 15.9 kDa. However, two larger, diffuse bands between 20-30 kDa were observed.

**Figure 2:**
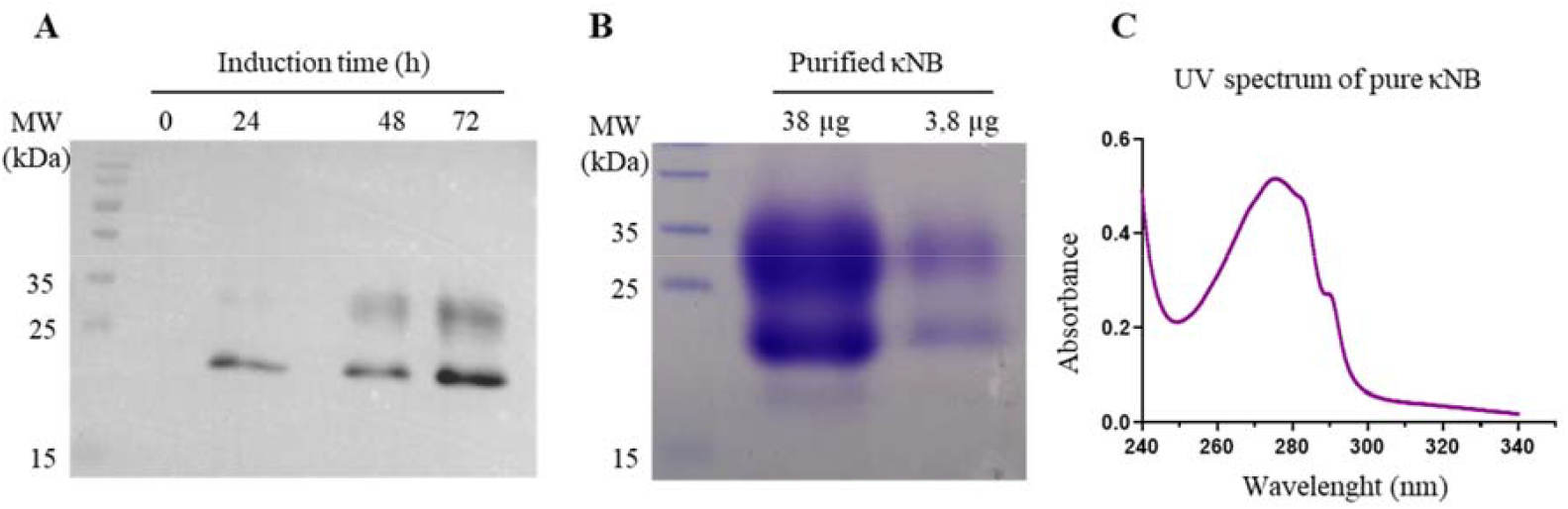
(A) κNB expression in *P. pastoris* induced supernatants. TCA-precipitated supernatants (300 µl) were run on a 15% SDS-PAGE, transferred to a PVDF membrane and incubated with mouse anti-His primary antibody (1:7500) and with an anti-mouse conjugated to HRP (1:15000) secondary antibody. The image obtained results from the superposition of the Western Blot with the image taken from the pre-stained marker (MW). Lanes to the right correspond to 0, 24, 48 and 72 hours of culture induction. **(B)** SDS-PAGE (15%) of pure κNB (loaded 38 µg and 3.8 µg of protein). The expected κNB size is 15.9 kDa. **(C)** UV-visible spectrum of a 3.8 µg of κNB after subtracting the buffer alone spectrum. The A280 nm value was 0.49.

Protein produced in 200 mL of supernatant of cultures induced during 72 hours was purified by affinity chromatography using a 1 mL Ni-NTA column. Bound κNB was eluted with 300 mM imidazole and fractions in which eluted NBs were identified were pooled and dialyzed against TBS. SDS-PAGE and staining with Coomassie Brilliant Blue after κNB purification and dialysis revealed the same two-band pattern previously shown by Western Blot (**Figure 2B**).

The UV spectrum of the pure protein corresponds to the expected profile considering the number of tryptophan and tyrosine residues in the sequence (**Figure 2C**). Protein concentration was quantified using the absorbance at 280 nm and the extinction coefficient of 32890 M^-1^cm^-1^, resulting in a pool of 2.37 mg/ml and, by extrapolation, a yield of 17.8 mg/L of culture.

The presence of the two diffuse bands of a lower mobility than the predicted molecular weight observed in **Figures 2A** and **2B** was attributed to the possible glycosylation of the Asn 107 located in CDR3 of the κNB during its transit through the yeast secretory pathway. Two κNB deglycosylation reactions were carried out to confirm this: one in native conditions and another one in denaturing conditions, using Endoglycosidase H (EndoH) to cleave high mannose glycans produced by yeast. The reactions were run in a new SDS-PAGE along with untreated controls (**Figure 3A**), showing that after treatment with EndoH the two diffuse bands become one concise and defined band of the expected size of 15.9 kDa. Based on these results, it was concluded that the presence of the two-band pattern was due to the *N*-glycosylation of the Asn 107 of κNB. Although the prediction of the complex structure suggested that the presence of an *N*-glycan would not affect NB-*kappa* domain interaction we evaluated how κNB’s performance and/ or function would be affected by the presence of glycans. It is worth mentioning that a third, very faint band was visible in the controls, indicating that a small fraction of the original κNB may have remained non-glycosylated and that *P. pastoris* may produce different glycosylation patterns in the same protein.

**Figure 3:**
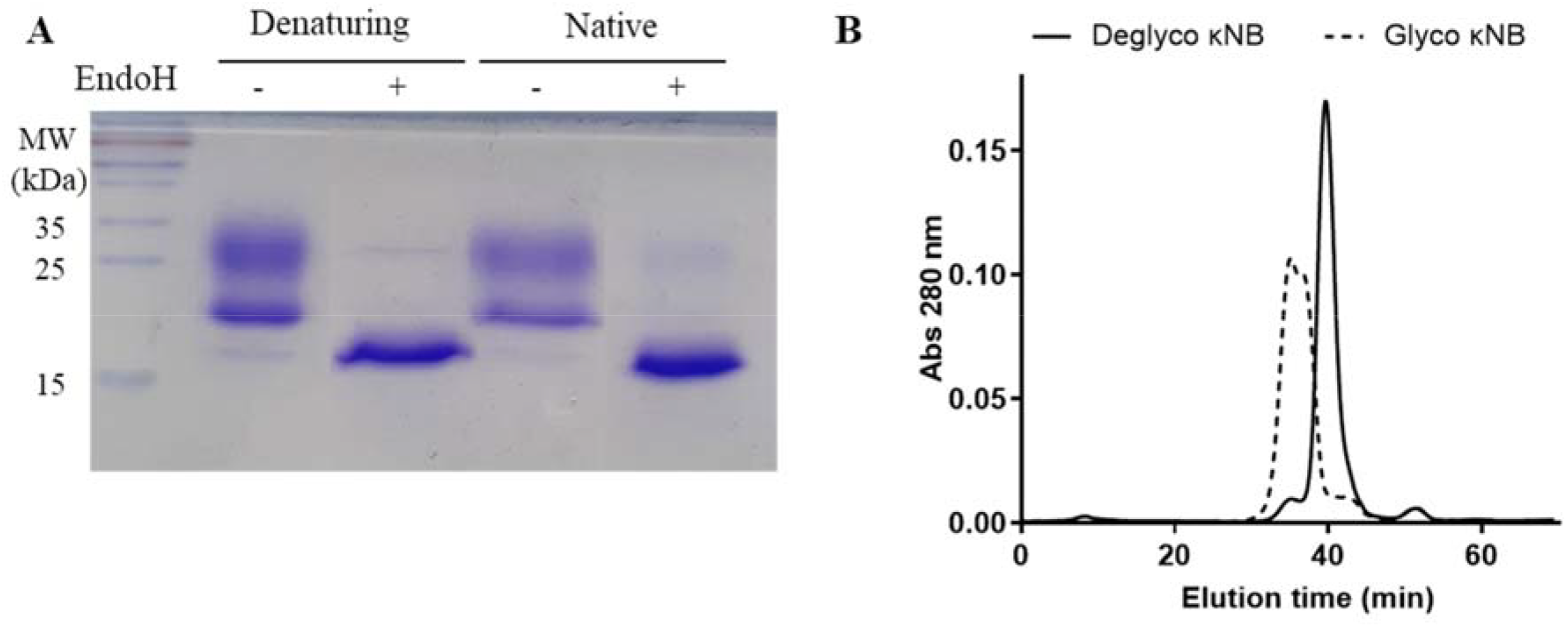
(A) Analysis of the glycosylation status of κNB expressed in *P. pastoris*. κNB was deglycosylated with EndoH under both denaturing and native conditions and analyzed on 15% SDS-PAGE. **(B)** SEC-HPLC profile of glycosylated κNB and κNB deglycosylated under native conditions. The graph represents the elution chromatogram of the different molecules, plotted as absorbance at 280 nm vs. elution time. Each sample was run for 70 minutes in 20 mM Tris-HCl, 100 mM NaCl, 1 mM EDTA, pH 7.0.

SEC-HPLC was carried out on both glycosylated and deglycosylated κNB in native conditions to evaluate purity, size and aggregation. The elution profile of deglycosylated κNB (**Figure 3B)** displayed a well-defined, narrow elution curve, with a peak at 40 minutes. On the other hand, the SEC profile of glycosylated κNB showed an earlier elution peak at 35 minutes, consistent with the presence of a larger molecule. Moreover, a wider peak was obtained for glycosylated κNB likely due to the heterogeneous glycosylation profile revealed both by SDS-PAGE and Western Blot.

### κNB functional tests

The functionality and specificity of the anti-*kappa* NB to recognize mouse antibodies was assessed by ELISA. A plate was coated in duplicate with serial dilutions of human, llama and mouse sera, followed by incubation with serial dilutions of both glycosylated and deglycosylated κNB to assess whether the presence of glycans in the κNB affects the antibody recognition. The κNB binding was detected with an anti-His-HRP antibody. Resulting data of absorbance at 450 nm showed that the κNB specifically recognized only mouse serum, without showing any detectable cross reactions with sera from human or llama origins, indicating that the κNB is highly species-specific (**Figure 4A & 4B**).

**Figure 4.**
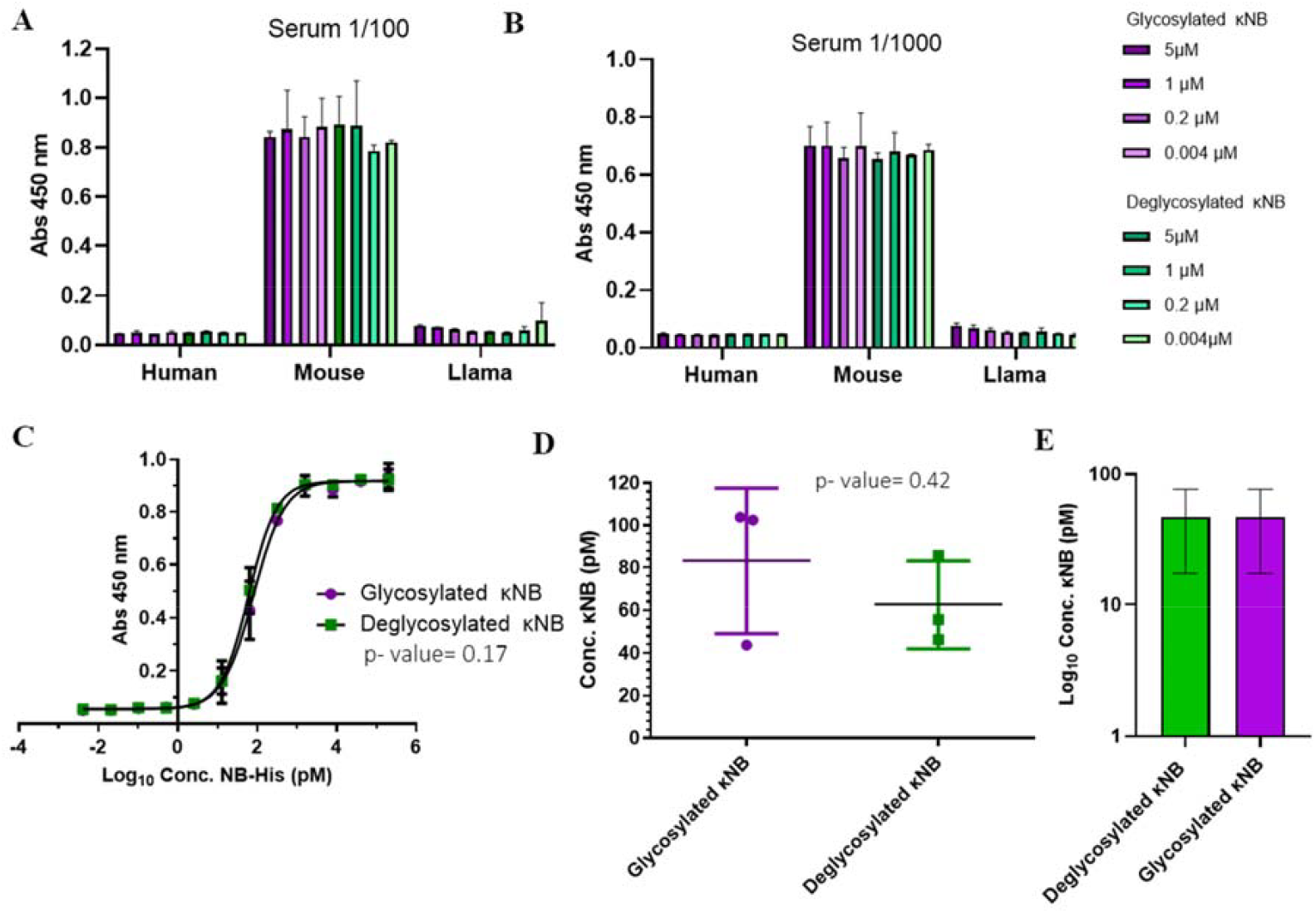
κNB specificity towards mouse antibodies. **(A)** Graph depicting κNB binding profile to an ELISA plate coated with human, mouse and llama sera in 1:100 dilution. **(B)** Graph depicting κNB binding profile to an ELISA plate coated in human, mouse and llama sera in 1:1000 dilution. Absorbance of TMB substrate was measured at 450 nm. **(C)** κNB detection limit of 1:1000 dilution of mouse serum. Absorbance was measured at 450 nm. Log_10_ of κNB concentrations were used a better visualization. **(D)** Comparison of EC50 of glycosylated and deglycosylated κNB. Standard deviations and p-value determined after a t-test are indicated. **(E)** Comparison of limits of detection determined for both glycosylated and deglycosylated κNB. The limit of detection was defined as the highest concentration of κNB that produces a signal at 450 nm that is greater than the cut off value set at x3 times the blank. Log_10_ of κNB concentrations were used a better visualization

Moreover, glycosylation did not affect κNB functionality or specificity, as glycosylated and deglycosylated κNB behaved indistinguishable one from another.

To determine κNB detection limits, a new ELISA was carried out using only 1:1000 mouse serum dilution and incubating with 200 nM to 4 fM serial dilutions of glycosylated and deglycosylated κNB. The interaction curve obtained after absorbance measurement showed that both forms of the κNB displayed identical performances (**Figure 4C**). To determine whether the differences between κNBs were statistically significant, an analysis of variance (ANOVA) was carried out defining a significance level (α) of 5%. The resulting p value of 0.17 confirmed that there are no significant differences between the glycosylated and deglycosylated κNBs and therefore κNB performance is not affected by the presence or absence of glycans, provided that they are removed *after* the protein synthesis.

The κNB’s EC50 (effective concentration, which was defined as the concentration at which 50% of the antibody is bound) were 80.9 pM for the glycosylated κNB and 61.2 pM for the deglycosylated κNB (difference determined not significant by a t-test, p-value 0.42) (**Figure 4D**).

The limits of detection were determined for both glycosylated and deglycosylated forms of the κNB. These were defined as the highest κNB concentration that gives a signal at 450 nm higher than the cutoff value set at X3 times the blank of the reaction. Using this criterion, the limit of detection was of 46.9 pM for both forms of κNB (**Figure 4E**).

### κNB-HRP expression in *P. pastoris* and functional tests

Once the smallest, simplest variant of the κNB expressed in *P. pastoris* proved to be specific, functional and sensitive, *P. pastoris* was transformed with the plasmid that would allow the expression of the κNB fused to HRP for chemiluminescent detection, and a Hisx6 tag for purification (κNB-HRP). Integration of the gene in the genome was confirmed by PCR, and induction of several clones was carried out in 10 ml cultures to quickly assess HRP enzyme activity in cultures supernatants. The better clone that displayed HRP activity measured by TMB colorimetric test was chosen for a 200 mL induction. κNB-HRP was purified as described for κNB using a Ni-NTA affinity column. Quantification by absorbance of the eluted and dialyzed pool was measured at 280 nm using an extinction coefficient of 45840 M^-1^cm^-1^. A concentration of 759.8 µg /mL was obtained for the purified pool resulting in a total yield of 7.2 mg κNB-HRP per liter of total culture.

The same as occurred when expressing κNB, κNB-HRP also presented a highly diffuse and heterogeneous pattern ranging from 60 to 100 kDa by SDS-PAGE that was sharpened to a single band of the expected size of 49 kDa after deglycosylation with EndoH **(Figure 5A)**. A few smaller bands than expected were observed upon deglycosylation, probably due to some extent of protein degradation. Western Blot using anti-His antibody confirmed κNB identity and that those smaller bands were degraded fractions of the κNB-HRP **(Figure 5B)**.

**Figure 5.**
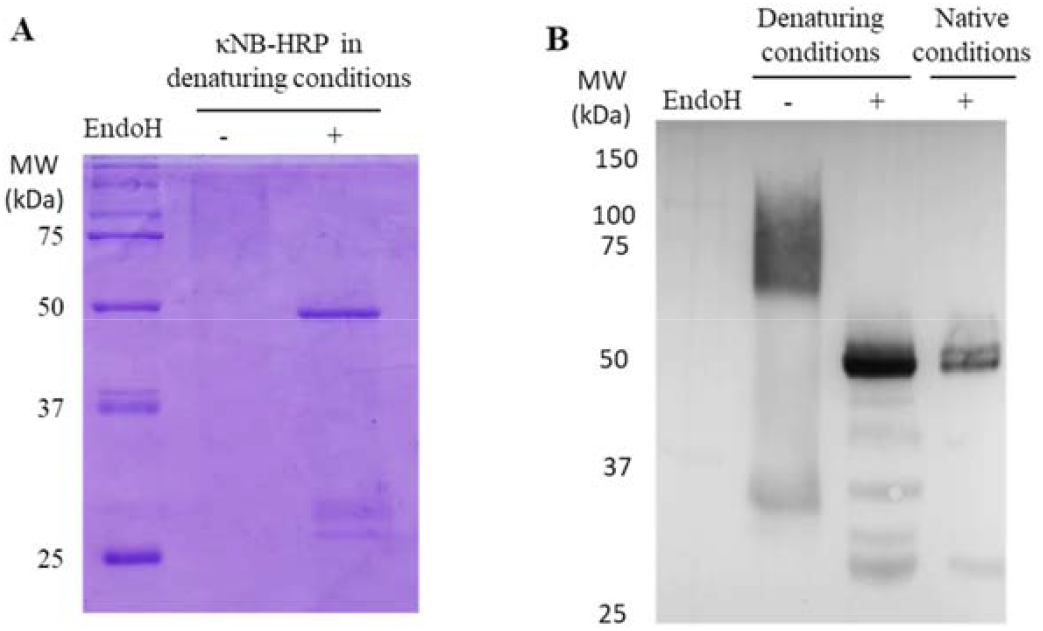
Expression of κNB-HRP in *P. pastoris*. **(A)** SDS-PAGE of pure κNB-HRP (5 μg) before and after treatment with EndoH. **(B)** Western Blot performed on κNB-HRP (5 μg) after deglycosylation in native and in denaturing conditions. Untreated controls were included. Mouse anti-His was used as primary antibody (1:7500) and anti-mouse-HRP as secondary antibody (1:15000).

An ELISA was carried out to test the κNB-HRP functionality. The plate was coated with both mouse serum, for a direct assay with κNB-HRP, and with SARS-CoV2 RBD protein for an indirect assay using mouse anti-RBD serum^[15]^ as primary antibody and κNB-HRP as secondary antibody. κNB-HRP was tested in the range 36 – 0.5 nM. In both assays, both glycosylated and deglycosylated κNB-HRP produced measurable signals at 450 nm, with the deglycosylated molecule producing higher average signal values (**Figure 6A & 6B**). The analysis of variances produced a p value lower than that of the α set at 5% for both tested conditions (p 1,8376E-07 for the assay against mouse serum and 1,9342E-11 against RBD and mouse anti-RBD serum), therefore in this case the difference in performance between the glycosylated and deglycosylated κNB-HRP is statistically significant, indicating that glycan presence in both κNB and HRP domains not only did not affect the function but in some cases improved the performance.

**Figure 6.**
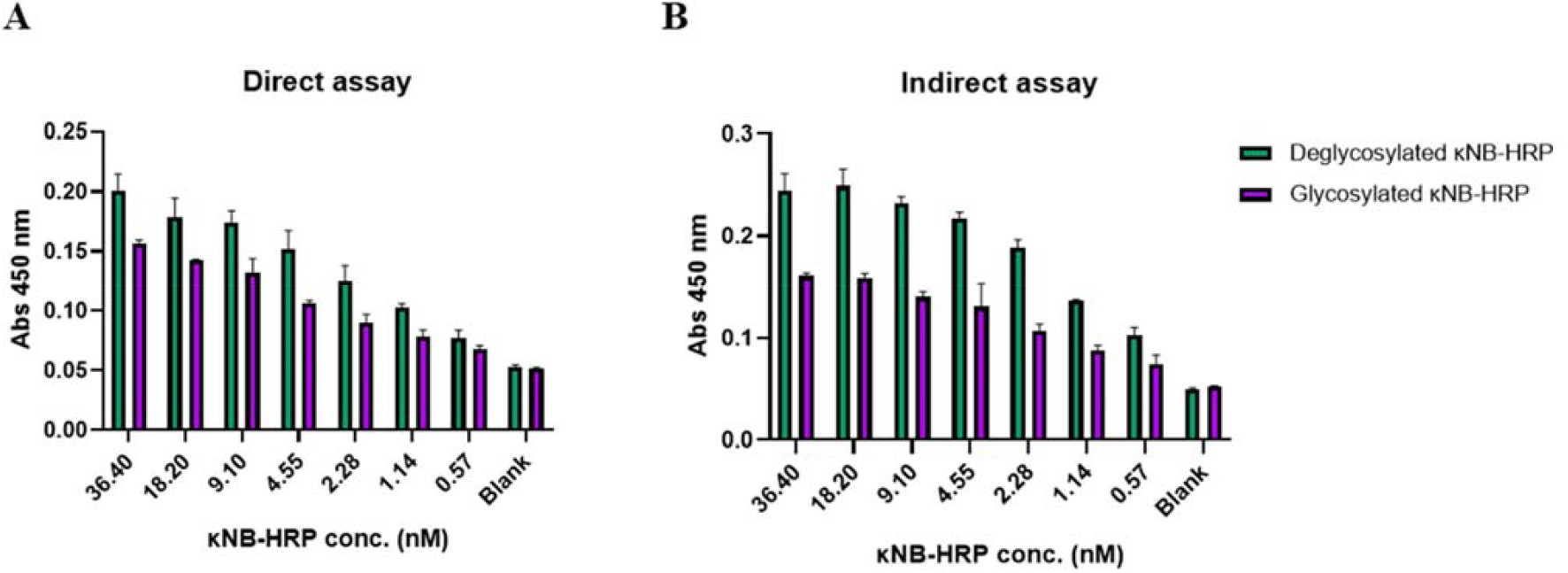
Interaction profile by ELISA of κNB-HRP with **(A)** Mouse serum in a direct assay **(B)** RBD protein and anti-RBD mouse serum in an indirect assay. κNB-HRP was tested in concentrations ranging from 36,4 to 0,5 nM and absorbance was measured at 450 nm.

κNB-HRP’s performance as a secondary antibody was also tested in a Western Blot. Different amounts of a total protein extract of the yeast *Schizosaccharomyces pombe* expressing a GFP-glucosidase I fusion protein were run in a polyacrylamide gel and the blocked membrane was incubated with a mouse anti-GFP primary antibody (1:1000). The membrane was divided, one half was used as a methodological control and incubated with a commercial anti mouse-HRP monoclonal secondary antibody diluted 1:15000 (**Figure 7A**); the second membrane was incubated with the κNB-HRP, deglycosylated in native conditions, diluted 1:3000 (**Figure 7B**). Both membranes showed the expected size band of the fusion control protein of approximately 120 kDa, and could detect 5 µg of total protein. These results show that the κNB-HRP is suitable to be used as a secondary antibody both in Western Blots and ELISAs.

**Figure 7.**
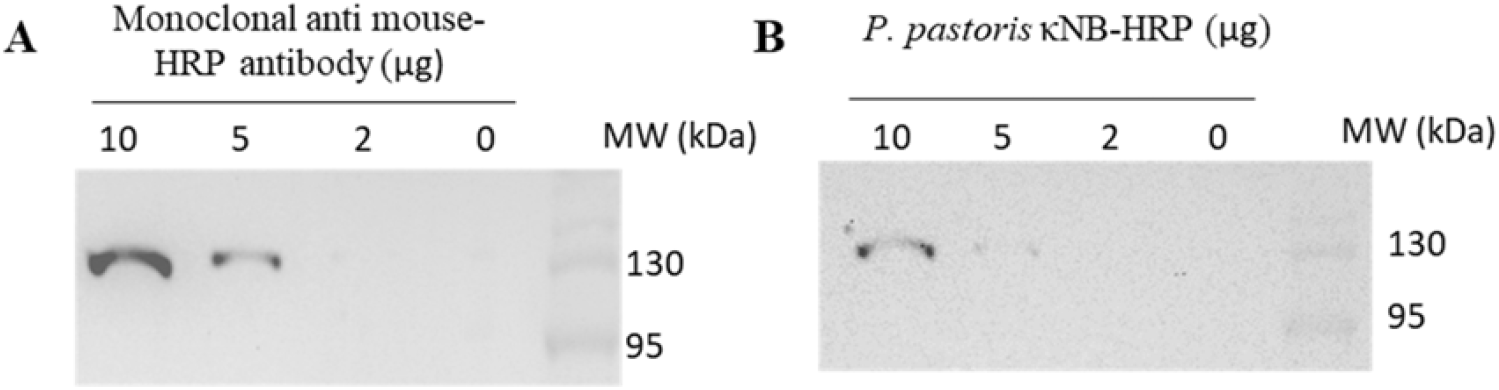
Use of κNB-HRP as a secondary antibody in a Western Blot. **(A)** an anti-mouse-HRP of known performance (1:15000) and **(B)** deglycosylated κNB-HRP in a 1:3000 dilution were used. Samples from 10 through 0 µg of total protein extracts expressing a control GFP fusion protein. In both cases, membranes were incubated with mouse anti-GFP (1:1000) as the primary antibody.

## DISCUSSION

The discovery of NBs 30 years ago revolutionized antibody engineering due to their structural characteristics and the advantages these confer over conventional antibodies. These benefits have proven useful for the development of potential applications in therapeutic, diagnostic, and research areas. Although expression of NB is popular in bacterial systems, expression in eukaryotic systems would allow post-translational modifications of both NBs and fused proteins or tags that may require glycans to fold properly.

In this work, a modular plasmid was designed for the expression in *P. pastoris* of an anti-mouse *kappa* chain NB fused to various accessory peptides of choice, which could be used in immunoassays as a secondary antibody, for immunoprecipitations, or for antigen capture in ELISA tests, with tags to facilitate NB purification and/or detection.

The Alpha fold prediction shows that the κNB-*kappa* light chhain complex is compact and well-structured, with a large CDR3 stretch of κNB being involved in the direct interaction. Moreover, we predicted that the presence of a glycan in the CDR3 would not affect the interaction surface. Further work will be done to investigate the internal motions of the CDR3 stretch to evaluate whether binding depends on the local dynamics of this region.

The modular vector was cloned into *E. coli* and three derived vectors were obtained: one carrying the whole modular plasmid, one for the expression of κNB and one for the expression of κNB-HRP. κNB and κNB-HRP constructions were integrated in the genome of *P. pastoris* and NBs expressed under the AOX1 methanol-induced promoter.

Anti-*kappa* mouse NB was purified in its monomeric form from the culture supernatant by Ni-NTA affinity column and a yield of 17.8 mg/L was obtained, one of the highest values reported for the expression of NBs in yeast^[16,17]^. The κNB was indeed glycosylated, a fact confirmed through treatment of pure protein with endoglycosidases. The glycosylated and non-glycosylated preparations were analyzed by Western Blot and SEC-HPLC, confirming that the size heterogeneity was due to the glycosylation in the eukaryotic system.

The specificity and functionality of glycosylated and non-glycosylated κNB were tested by ELISA. These assays demonstrated the specificity of both κNBs, which interacted only with mouse serum and not with that obtained from other species tested. The detection limits, EC50 values, and variance analysis revealed no statistically significant differences in performance between the glycosylated and non-glycosylated NBs (with a p-value of 0.17), indicating that, as predicted, the presence of *N*-glycans did not affect its functionality.

*N*-glycosylation is involved in the quality control of glycoprotein folding in the endoplasmic reticulum^[18]^. The presence of a glycan in the molecule in that compartment favors that only properly folded glycoproteins follow the secretory pathway of eukaryotic cells and thus are secreted to the culture media, otherwise they are retrotranslocated to the cytosol from the endoplasmic reticulum and degraded. This ensures the native folding status of proteins obtained from culture’s supernatants. Previous work from our group has shown that there were no structural differences between glycosylated and deglycosylated Spike’s receptor binding domain (RBD)^[15]^. Deglycosylation showed a more homogeneous profile, indicating that glycan could be beneficial for obtaining a higher yield of folded expressed proteins, and that heterogeneity produced could be removed afterwards if needed for regulation purposes.

The second NB variant induced and purified from the modular construction was κNB-HRP, to produce a useful secondary antibody replacement. The protein was obtained at a calculated yield of 7,2 mg of protein/L of culture in shaked flasks. κNB-HRP proved to be functional after successful trials in both ELISAs and Western Blot. κNB-HRP was also glycosylated in both κNB and HRP, but in this case deglycosylation showed a slightly better performance that the glycosylated counterpart. It is worth mentioning that while HRP has been expressed in prokaryotic systems, such as *E. coli*, several difficulties have been reported: affected stability due to lack of glycosylation, low refolding efficiency of inclusion bodies, and reduced catalytic activity after refolding^[19]^; all of which positions *P. pastoris* as a great tool to express different conjugated NBs.

As proof of concept, the vector was used to produce κNBs either alone or fused to HRP. However, we produced a highly useful tool with the potential for broader applications in the future-such as improving the orientation and immobilization of NBs on the polystyrene surfaces of ELISA plates or enabling biotinylation-based modifications that could expand the range of κNB applications. Also, it is important to mention that the high cost and significant delays associated with importing research antibodies hinder access to these crucial reagents in many low-income countries. Establishing a local, efficient NB production system could offer a valuable solution to reduce costs and wait times, fostering local research and development.

## EXPERIMENTAL PROCEDURES

### Materials

Yeast extract, tryptone and agar were purchased from Britania (Argentina). Yeast nitrogen base and peptone were from Difco. Zeocin, Taq polymerase and dNTPs were from Invitrogen. Sorbitol, amino acids for culture media and RNAse A were from Sigma. Dextrose was from Biopack. DNA restriction enzymes and bovine seroalbumine (BSA) were from New England Biolabs. T4 DNA Ligase was from Promega. Oligonucleotides were synthesized by GenScript. *SuperSignal® West Pico Chemiluminescent Substrate* for Western Blot and *SnakeSkin™ Dialisys Tubing* 10.000 MWCO for protein dialysis were acquired from ThermoFisher Scientific.

### Strains and Media

*Escherichia coli* DH5α strain was used for cloning and amplification purposes. *E. coli* growth was carried out at 37°C in low salt LB medium (Luria-Bertani broth; 1% tryptone, 0.5% yeast extract, 0.5% NaCl, pH 7.5). For the selection of transformed bacteria, zeocin was added up to a concentration of 25 µg/ml. *P. pastoris* wild type strain X-33 (Invitrogen) was used for protein expression. Yeasts were grown in YPDA medium (Yeast extract Peptone Dextrose Adenine medium;1% yeast extract, 2% peptone, 2% glucose, 75 mg/L adenine). Transformed yeast cells were selected using YPDS medium with zeocin (Yeast extract Peptone Dextrose Sorbitol medium; YPDA supplemented with 1M sorbitol and 100 µg/mL zeocin). For protein expression, cells were first grown in BMGY medium (Buffered Glycerol complex Medium; 1% yeast extract, 2% peptone, 100 mM potassium phosphate buffer pH 6.0, 1% glycerol, 1.34% yeast nitrogen base, 4×10^−5^ % biotin). The protein expression was induced in BMMY medium (Buffered Methanol complex Medium; BMGY in which glycerol is replaced by 1% methanol as carbon source). Yeast cultures were grown at 28°C. In all cases, solid plates were prepared adding 2% agar.

### Prediction of κNB-mouse *kappa* light chain complex structure

The structure of the complex between the nanobody and the *kappa* domain (mouse) was predicted employing AlphaFold3 at https://alphafoldserver.com/ ^[20]^. The sequence corresponding to the *kappa* domain was obtained from the protein sequence database at NCBI. We used the sequence of the immunoglobulin *kappa* light chain variable region from *Mus musculus*. On the other hand, the sequence corresponding to the nanobody was found in reference Pleiner *et al*. (2018)^[13]^. Five different models of the complex were obtained and analyzed using standard parameters. For the analysis pTM and ipTM scores were considered. The predicted template modeling (pTM) score and the interface predicted template modeling (ipTM) score are both derived from the template modeling (TM) score that measures the accuracy of the entire structure^[21,22]^. A pTM score above 0.5 suggests that the predicted complex’s overall shape is probably similar to the real structure. The ipTM metric measures how accurately the relative positions of the subunits within the complex are predicted. Scores over 0.8 indicate reliable, high-quality predictions, while those below 0.6 point to possible prediction failure. ipTM values between 0.6 and 0.8 fall into a gray area where predictions may or may not be correct.

### Plasmid design

The individual coding sequences for a NB that recognizes the *kappa*-chain of mouse immunoglobulins, a plastic binding sequence, the HRP enzyme and the Avitag sequence were previously described by Pleiner *et al*. (2018)^[13]^, Qiang *et al*. (2017)^[23]^, Yamagata & Sanes (2018)^[24]^ and Athavankar & Peterson (2003)^[25]^, respectively. The sequences were codon optimized for expression in *P. pastoris* and designed to be incorporated into the pPICZαA vector (Invitrogen) using the molecular biology tool Benchling (https://www.benchling.com/). The sequences were placed in a particular order with different restriction sites which would allow the removal of each accessory peptide and subsequent religation to obtain different vectors able to express combinations of fusion proteins. The construction was then synthesized by GenScript and cloned complete into the pPICZαA vector, resulting in a new secretion vector named pPICZαA-Plastic-κNB-HRP-Hisx6-Avitag (also referred to as “Modular plasmid” in this work). Once synthesized, the vector was transformed into *E. coli* DH5α strain and extracted by minipreparation. *E. coli* DH5α strains containing three derived vectors were obtained: one carrying the whole modular plasmid, one carrying a plasmid for the expression of κNB (κNB expression fused to a Hisx6 tag; pPICZαA-κNB-Hisx6), and one for the expression of κNB-HRP (κNB in fusion to the HRP enzyme and a Hisx6 tag; pPICZαA-κNB-HRP-Hisx6).

### DNA procedures

*E. coli DH5*α chemocompetent cells were prepared as detailed in [26]. After recovery, cells were plated in LB low salt agar with zeocin 25 µg/mL and incubated overnight at 37°C. Miniprep DNA isolations from bacteria were carried out as detailed in [26]. When DNA was purified from 100 mL cultures, GenElute™ HP Plasmid Midiprep Kit (Sigma Aldrich) was used following the manufacturer’s instructions. DNA digestions with restriction enzymes were performed at 37°C overnight and complete digestion was confirmed by 1% agarose gel electrophoresis. Vectors without the removed module were extracted form gel using either the *Monarch® DNA Gel Extraction Kit* from New England Biolabs or *QIAquick® Gel Extraction Kit* from Qiagen. DNA re-ligation reactions were done adding T4 DNA ligase (Promega) and incubating overnight at 10°C, after which the ligase was heat-inactivated and the plasmid transformed into *E. coli* for recovery. The correct removal of sequences after each restriction was verified by both colony PCR and sequencing. For DNA quantifications after gel electrophoresis the samples were visualized using the InGenius3 gel documentation system and GeneSys software, and compared with a standard using the *GelAnalyzer* software (www.gelanalyzer.com) (Istvan Lazar Jr., PhD & Istvan Lazar Sr., PhD, CSc).

### *P. pastoris* genetics procedures

Electrocompetent *Pichia pastoris* cells were prepared as described in *[27]*. The desired plasmids bearing the combination of fusion proteins for expression were linearized using SacI to direct integration in *P. pastoris* AOX1 promoter. Transformation of yeast was performed with linearized plasmid DNA (>5 µg) by electroporation at 2.5 kV, 25 µF and 200 Ohm. Recovered cells were selected in YPDS plates with 100 µg/ml zeocin and incubated at 28°C for 5 days. Integration of the correct modular combination plasmid into *P. pastoris* genome was confirmed by colony PCR using the following primers: 5’ AOX1 (5’-GACTGGTTCCAATTGACAAGC-3’), 3’AOX1 (5’-GCAAATGGCATTCTGACATCC-3’), Plastic-rev (5’-CACCACTGTCTGAAATCCCA-3’), NB-rev (5’-GTCCGCATAGTATGTGTAGC-3’), HRP-rev (5’-TTGTCGTAGAAGGTAGGCGT-3’), *AviTag-rev (*5’-GATGTCGTTTAGGCCACTAG-3’).

### κNB induction and purification

For each κNB expression, a single colony was inoculated in a starter culture of 20 ml BMGY and grown at 28°C with 250 rpm agitation. Cells were then centrifuged, resuspended to OD=1 in BMMY and incubated at 28°C with agitation at 250 rpm for 72 hours, adding 1% methanol every 24 hours. After 72 hours, the culture was centrifuged at 3000 x g for 10 minutes and the supernatant was frozen until purification. κNB was purified from 200 ml culture supernatant using a 1 mL Ni-NTA affinity column and as described in ^[15]^. Fractions that contained protein eluted with 250 mM imidazole were identified by Bradford and run on a 10% Sodium Dodecyl Sulfate Polyacrylamide Gel Electrophoresis (SDS-PAGE). Fractions were pooled and purified protein was dialyzed against TBS (20 mM Tris-HCl, 150 mM NaCl, pH 7,4). Pure protein was quantified by UV spectroscopy, measuring absorbance at 280 nm using the following extinction coefficients: for κNB = 32890.00 M^-1^cm^-1^ and for κNB-HRP = 45840.00 M^-1^cm^-1^, assuming fully reduced cysteines. κNB size was verified by 10-15% SDS-PAGEs runs stained with Coomassie Brilliant Blue. Alternatively, to verify protein expression gels were transferred to PDVF membranes for Western Blotting during 80 minutes at 100 V in 20% methanol, 25 mM Tris, 192 mM glycine) buffer. Western Blot membranes were blocked with 3% low fat milk and incubated with a mouse anti-His (1:7500, Roche) as primary antibody and anti-mouse-HRP (1:15000) as secondary antibody, alternatively mouse anti-GFP (1:1000, Roche) was used when testing the NB as secondary antibody. Membranes were revealed using the *SuperSignal® West Pico Chemiluminescent Substrate* (ThermoFisher Scientific) and visualized in a GeneGnome imaging system using the GenSys software (10 minutes exposure with 1 minute intervals or 20 minutes exposure with 2 minute intervals).

### κNB deglycosylations

High-manose glycans were removed from pure κNB in native conditions using 14.4 mU of Endoglycosidase H (EndoH) produced in *P. pastoris* in house as described^[15]^. For glycan removal in denaturing conditions, κNB were first denatured by heating at 95°C for 10 minutes in denaturing buffer (0.5% SDS, 40 mM DTT) and then incubated with 1.44 mU of EndoH at 37°C for 1 hour. Analysis were performed either by SDS-PAGE or Western Blot.

### Size exclusion-high-performance liquid chromatography (SEC-HPLC)

Purified κNBs were injected into a Superose-6 column (GE Healthcare) coupled to a JASCO HPLC equipment with a UV-VIS UV-2075 detector. The running buffer composition was 20 mM Tris-HCl, 100 mM NaCl, 1 mM EDTA, pH 7,0. The experiment was run for 70 minutes at room temperature (∼ 25°C), with a flow set to 0.4 ml/min and elution was monitored at 280 nm.

### ELISA analysis

Flat-bottom 96-well ELISA plates (Greiner Bio-One) were coated with human, llama or mouse blood serum diluted in carbonate buffer (32 mM Na2CO3, 68 mM NaHCO3) and incubated overnight at 4°C. After serum adhesion, plates were washed 3 times with water and once with PBS-T (NaCl 0,14 M, KCl 2,7 mM, Na_2_HPO_4_ * 7 H_2_O 10 mM, K_2_HPO_4_ 1.4 mM, Tween 20 0,1%, pH 7.4). Blocking buffer (3% milk in PBS-T) was then added and incubated for 1 hour at room temperature. Further incubations were carried out in the same conditions and followed by the same washing steps. The NBs were diluted to the desired concentration in 1.5% low-fat dry milk in PBS-T and added to the wells. When testing the κNB against mouse serum, an anti-His tag-HRP antibody (HRP Anti-6X His tag® antibody, Abcam) was used in a 1:5000 dilution as secondary antibody. The κNB fused to HRP was tested in a direct assay in a plate sensitized with mouse serum (1:2400). For testing as secondary antibody in an indirect assay, the plate was coated using SARS-CoV2 receptor binding domain (RBD) and anti-RBD mouse serum was used as primary antibody in a 1:300 dilution. A commercial anti-mouse-HRP antibody was used as control secondary antibody (Anti-Mouse IgG (H+ L) HRP conjugate, Promega, 1µg/µL) in serial dilutions (ranging 1:250 to 1:16000). After the final incubation and washing, TMB substrate (*TMB Substrate Reagent Set, BD Biosciences)* was added and the absorbance at 450 nm was measured using an Infinite M Plex plate reader (Tecan).

### HRP fast Activity Test

To assess the activity of the HRP enzyme expressed in *P. pastoris* expressing κNB-HRP, 10 μL of 1/100 dilutions of supernatants from the induced culture medium supernatant were mixed with 5 μL of TMB substrate (*TMB Substrate Reagent Set, BD Biosciences)*. A positive result was rapidly visualized by a color change from clear to blue (progressing to brown upon saturation).

### Statistical analysis and computational work

Analysis of variance (ANOVA) was performed with Two-Factor with replication, defining an α of 0.05 (5%) in Microsoft Excel. Detection limit or end-point titer was defined as the NB concentration that produced a signal at 450 nm that was greater than the cutoff value, which was set at 3 times the average blank value. Calculation of the antibody EC50, was done using GraphPad Prism 5 software. Nonlinear regression (curve fit) and Sigmoideal dose-response (variable slope) models were selected.

## ACNOWLEDGEMENTS

This work was supported by grants from the Agencia Nacional de Promoción de la Investigación, el Desarrollo Tecnológico y la Innovación (ANPCyT) PICT2020-3099 and PICT2019-0016. Salary of researchers was supported by Consejo Nacional de Investigaciones Científicas y Técnicas (CONICET) and Universidad de Buenos Aires (UBA). SO is a fellow from ANPCyT. The *P. pastoris* strain used for expression of EndoH was the same used in Idrovo *et al.* 2024^[15]^, sent by Lixin Ma (College of Life Sciences, Hubei University, Wuhan, People’s Republic of China).

## Notes

### Competing Interest Statement

The authors have declared no competing interest.

